# LPATH: A semi-automated Python tool for clustering molecular pathways

**DOI:** 10.1101/2023.08.17.553774

**Authors:** Anthony T. Bogetti, Jeremy M. G. Leung, Lillian T. Chong

## Abstract

The pathways by which a molecular process transitions to a target state are highly sought-after as direct views of a transition mechanism. While great strides have been made in the physics-based simulation of such pathways, the analysis of these pathways can be a major challenge due to their diversity and variable lengths. Here we present the LPATH Python tool, which implements a semi-automated method for linguistics-assisted clustering of pathways into distinct classes (or routes). This method involves three steps: 1) discretizing the configurational space into key states, 2) extracting a text-string sequence of key visited states for each pathway, and 3) pairwise matching of pathways based on a text-string similarity score. To circumvent the prohibitive memory requirements of the first step, we have implemented a general two-stage method for clustering conformational states that exploits machine learning. LPATH is primarily designed for use with the WESTPA software for weighted ensemble simulations; however, the tool can also be applied to conventional simulations. As demonstrated for the C7_eq_ to C7_ax_ conformational transition of alanine dipeptide, LPATH provides physically reasonable classes of pathways and corresponding probabilities.

**TOC Graphic:** 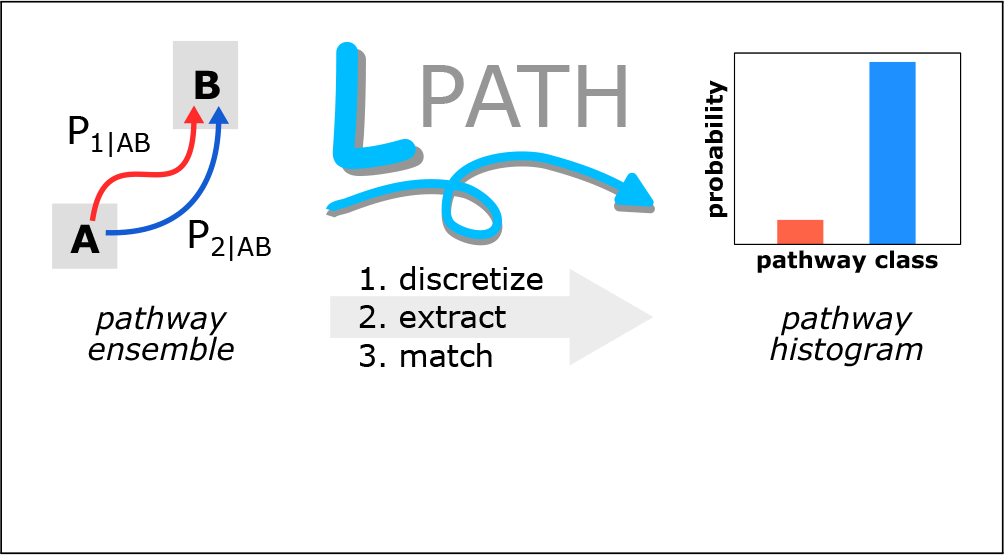

## Introduction

Pathways traversed by a molecular process–including all stable and transient states–are the most direct views of the mechanism by which the process occurs. Recent advances in both methods and hardware for physics-based molecular simulations have enabled the generation of these direct views for ever more complex processes that are beyond the reach of typical computing resources. Path sampling strategies–designed to focus sampling on transition pathways^1^ –have captured pathways (and rates) for processes such as chemical reactions, ^2,3^ crystal nucleation,^4^ binding processes of proteins^5,6^ and DNA,^7^ and large-scale conformational switching in proteins, ^8,9^ with orders of magnitude greater efficiency than conventional molecular dynamics (cMD) simulations. Furthermore, state-of-the-art supercomputers and dynamics engines have enabled simulations to target million-atom systems. ^10–12^

As we begin this golden age of molecular simulation, a next frontier is to gain a detailed understanding of how key processes are impacted by the multiple pathway routes that may exist. Identifying pathway routes, however, can be a challenge due to two factors. First, the clustering of molecular pathways into distinct routes can be non-trivial due to the large diversity and variable lengths of the pathways. Second, pathway analysis can be computationally intensive for complex processes due to the massive amount of simulation data generated (e.g., tens of TB^5,9^).

Current methods for pathway analysis involve two main steps: (1) projecting pathways onto a low-dimensional configuration space and (2) clustering pathways based on a similarity score. For example, the pathway similarity analysis (PSA) method^13^ is a “bottom-up” approach that projects pathways onto a low-dimensional configuration space consisting of the pairwise root-mean-squared deviation of sampled conformations and then clusters the pathways based on pairwise Hausdorff^14^ or Fréchet^15^ geometric distances. Another example is the pathway histogram analysis of trajectories (PHAT) method,^16^ which presents two approaches to classifying trajectories: (i) a “bottom-up” approach in which “set similarities” are used to generate similarity scores between pathways (e.g., using the geometric distances used in PSA) followed by Voronoi clustering of the pathways, and (ii) a “top-down” approach where fundamental sequences are calculated from a Markov state model (MSM). Most recently, MSMs have been used to train deep-learning models for latent-space path clustering (LPC).^17^

Here, we present the Linguistics Pathway Analysis of Trajectories with Hierarchical clustering (LPATH) tool, which clusters pathways using a bottom-up approach and a similarity score inspired by the Gestalt pattern matching algorithm for plagiarism detection.^18^ We adapt this score for the context of molecular pathways of variable lengths. Our projection of pathways onto one-dimensional text strings greatly accelerates the clustering of pathways and subsequent analysis of path ensembles relative to manual analysis of individual pathways. While the LPATH tool is designed for weighted ensemble (WE) path sampling, ^19,20^ using the WESTPA software package,^21^ this tool can also be applied to cMD simulations. Our benchmark application of LPATH involves the C7_eq_ to C7_ax_ conformational transition of alanine dipeptide. While alanine dipeptide is a relatively simple system that can be extensively sampled, it is sufficient in complexity for the purposes of this application note, which is to demonstrate the features of our software (i.e. the LPATH tool) for a benchmark system in which the validity of the resulting pathway classes can be clearly evaluated.

### The LPATH workflow

The LPATH workflow involves three steps (Figure 1): 1) discretization, 2) extraction, and 3) matching. An additional plotting module `lpath.plot` is available for visualizing LPATH results (i.e. pathway class histograms, event duration distributions, directed network plots) but not discussed here.

**Figure 1.**
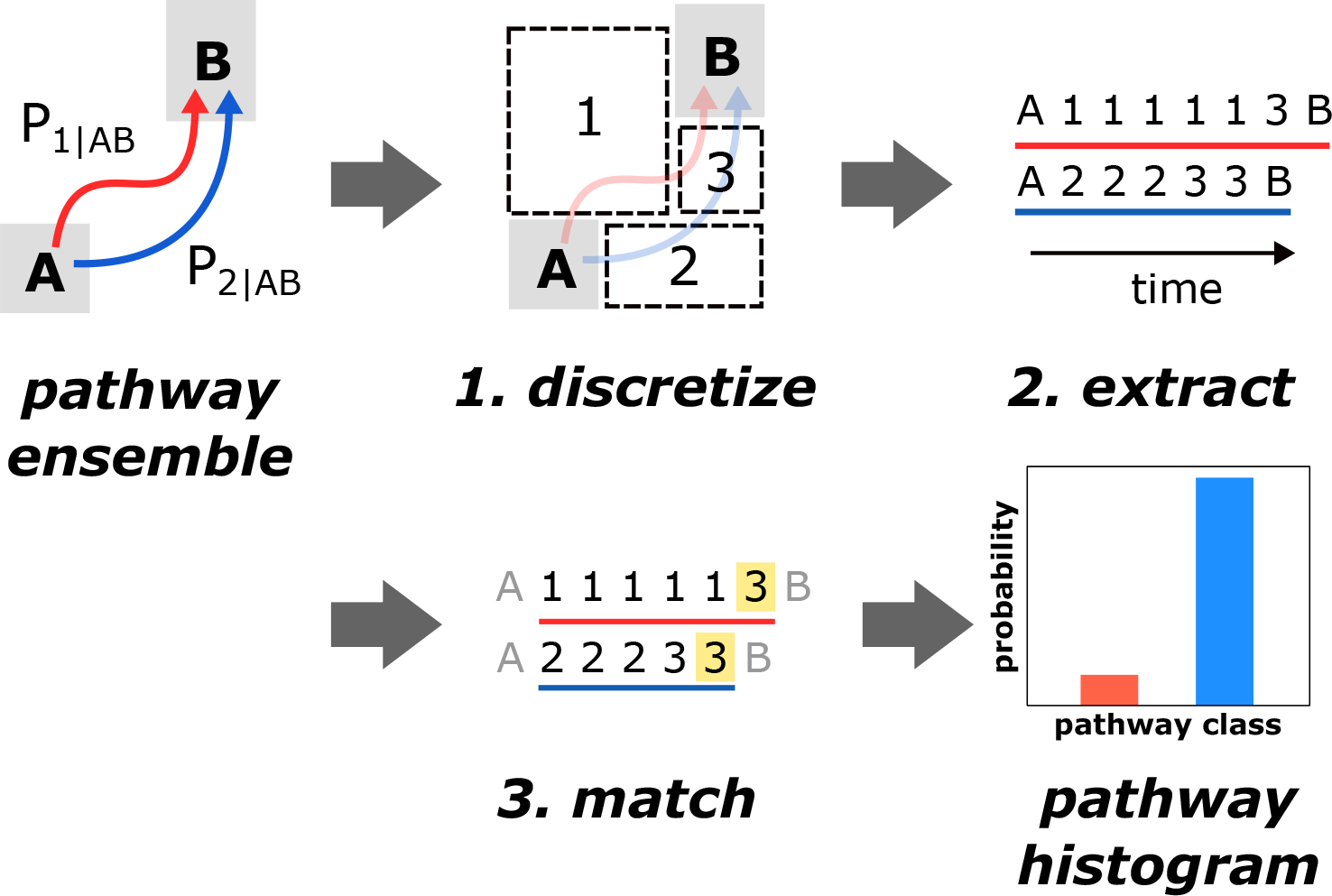
LPATH workflow for clustering pathways. The workflow consists of three steps executed at the command line. In the “discretize” step, source and target states are defined, and the configuration space between the source and target states is subdivided into discrete states. In the “extract” step, each successful pathway is represented as a text string consisting of the sequence of states visited at a specified frame frequency. The final “match” step calculates a similarity score between each pair of pathway text strings, then uses these similarity scores to perform hierarchical clustering of the pathways. The end result is a histogram of distinct pathway classes.

### Step 1: Discretize the configuration space

In this step, source and target states are defined, and regions of configuration space between these states are subdivided into discrete states. For a WE simulation, the ‘lpath.discretize’ module uses WESTPA’s `w_assign` tool to assign trajectory segments to the source and target states according to a scheme defined by the user in the `west.cfg` file. State discretization in the `west.cfg` file relies on first defining rectilinear bin boundaries as a list and then defining states as points, where the bin a point falls into becomes that state. The resulting `assign.h5` file is then used in step 2 (extraction step) to identify successful pathways connecting the source and target states. For a cMD simulation, trajectories can be assigned to source and target states with a Python function specified by the `--assign-function` option of `lpath.discretize`. The resulting `.npy` file is then used in the pathway extraction step.

Several options are available for discretizing the configuration space between the source and target states. Clustering methods, such as k-means clustering, can be used to assign states for both WE and cMD simulation datasets by supplying a user-defined Python function to the `--assign-function` option of `lpath.discretize` and output cluster labels as a `.npy` file. For WE simulation datasets, the unique number identifiers of trajectory segments (WE segment IDs) at each iteration can be used as proxy “states”. The use of trajectory segment IDs for discretization is beneficial in cases where configuration space is difficult to discretize based on one or two features. However, pathways among replicate WE simulations cannot be directly compared using this method, as segment IDs are not directly comparable between different simulations. If not using clusters or segment IDs as states, the configuration space of WE datasets can be discretized using WESTPA’s `w_assign` tool.

### Step 2: Extract successful pathways

This step extracts all successful pathways that connect the source and target states and saves these pathways as a convenient text-string sequence of states visited, along with other relevant simulation data (e.g., trajectory weights, progress coordinates, other properties for defining states). Options for fine-tuning this extraction step are described below.

First, we recommend using the finest time resolution (i.e. “frame” frequency at which conformations are captured) relevant to the completed simulation, which will be highly system-dependent. The choice of time resolution can have a major impact on the classification of pathways, i.e. pathways with a relatively coarse time resolution may miss key state-to-state transitions. The time resolution is specified using the `--stride` option and operates differently for WE vs cMD simulations. By default, analysis of WE simulations is done on frames every WE resampling time interval *τ*. However, users may set the `--stride` option to consider frames at a sub-*τ* time resolution, e.g. `--stride=10` for a WE simulation with *τ* =100 ps and frames saved every ps specifies frames every 10 ps. The resulting pathways would be 10x longer than those using frames every *τ*. If a pathway exits the source state or enters the target state in the middle of a *τ* interval, only the sub-*τ* frames after the exit or before the entry are considered. For cMD simulations, a stride of 10 provides 10x fewer points than the default resolution, considering conformations every 10 frames instead. The `--stride` option is a shared parameter that can be used in both the discretize and extract steps.

Second, we recommend removing shorter pathways (e.g., fewer than 10 sequential frames) that can exhibit inflated similarity scores by specifying a pathway length threshold using the `--exclude-length` option. The LPATH tool will automatically alert users if pathways with fewer than 10 frames exist in the pathway ensemble. Users may also consider higher thresholds when the pathway ensemble consists of more than 25% pathways below the default 10 frame threshold. However, users should ensure that each pathway class identified contains at least 10 pathways for a statistically-robust analysis.

### Step 3: Match pathways

This step calculates pairwise pathway similarity scores and identifies distinct pathway classes using hierarchical agglomerative (bottom-up) clustering. Any user-defined pathway similarity function written in Python can be used for this step with the `--match-metric` option. The output of the pathway similarity function should return a number that indicates the relative similarity of each pair of pathway strings. All pairwise similarities are then compiled into a distance matrix and clustered with the hierarchical agglomerative (“bottom-up”) clustering approach using the Ward linkage method.^22^

When matching, repeating patterns of states can often prevent a clear separation of pathways into distinct classes. To address this issue, we provide an option to condense these repeating patterns of user-specified lengths. For example, a `--condense 2` option will sequentially eliminate consecutive repeating characters (e.g. 11221112221122 becomes 121212) and consecutively-repeated pairs (e.g. 121212 becomes 12) to provide a fundamental sequence of states that disregards the length of time spent in each state. The ability to condense pathway strings greatly reduces the effects of pathway length on matching by focusing the pattern matching on the fundamental sequence of states visited, thereby improving the ability of the clustering algorithm to generate distinct pathway classes.

### A modified Gestalt pattern matching algorithm

The LPATH similarity score for a pair of pathways A and B is based on the Gestalt pattern matching algorithm, which is commonly used for plagiarism detection in computational linguistics:^18^

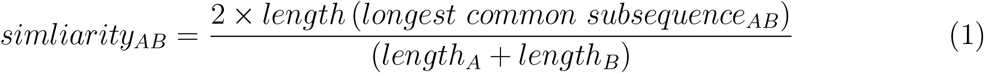

where the length of the longest common subsequence (see Figure 2A) is multiplied by two (to account for the fact that a pair is being evaluated) and divided by the combined length of the two pathway text strings being compared. The division by the combined lengths effectively normalizes the numerator (double the longest common subsequence) and provides a similarity score out of a maximum value of one.

**Figure 2.**
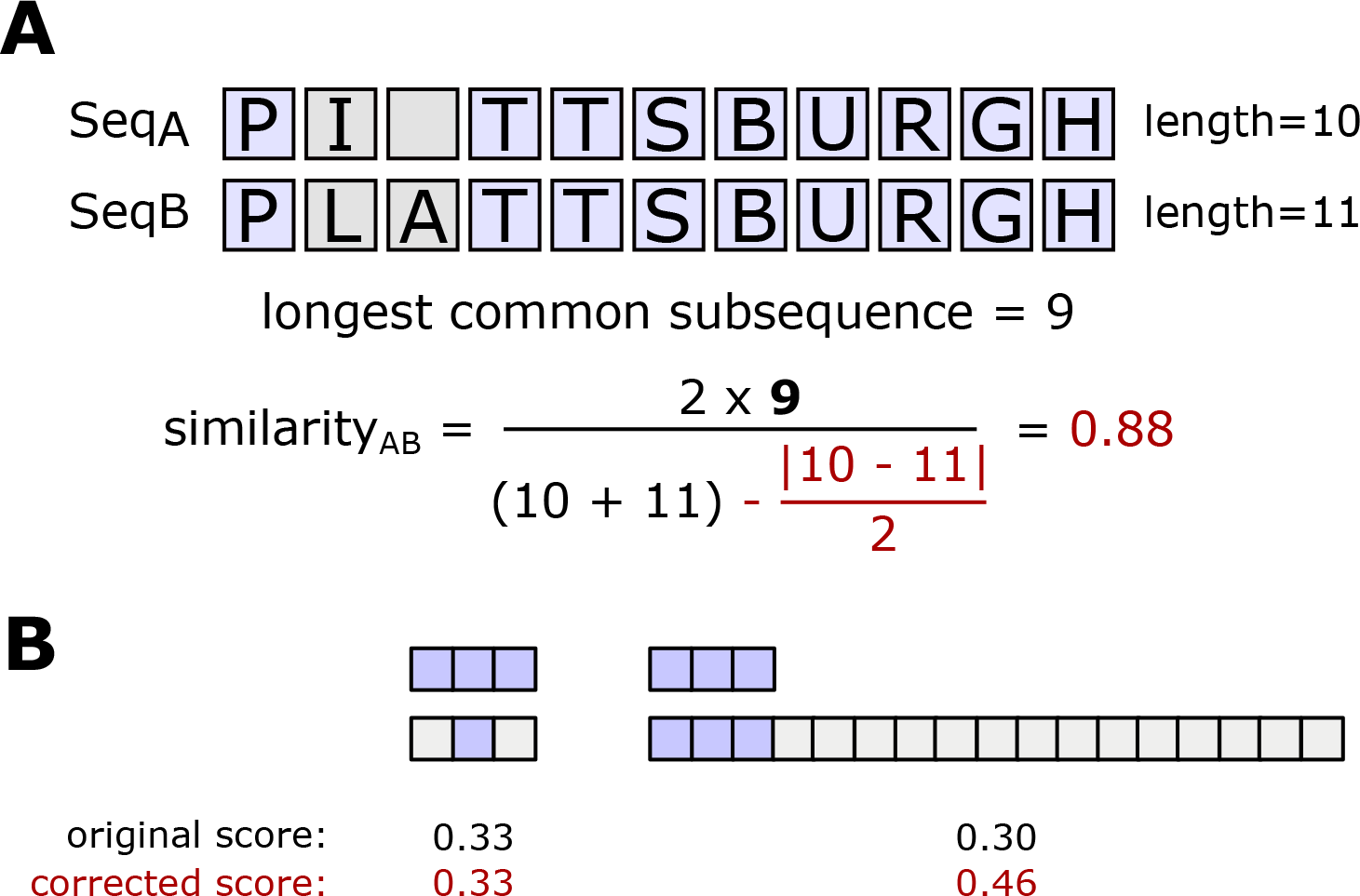
Illustration of the Gestalt pattern matching algorithm with a correction term for the comparison of molecular pathways. A) An example comparing two text strings with our corrected Gestalt pattern matching algorithm. These strings have a longest common subsequence of nine characters and lengths of ten and eleven, respectively. The assessed similarity between these strings is 0.88, an expected result given how similar these two strings appear. B) An example of how the original Gestalt pattern matching algorithm can be inaccurate when comparing string pairs of dramatically varying lengths. Each colored block represents one of two characters, purple or gray. The main string in this example (three purple) appears more similar to a string of the same length (gray-purple-gray) than it does to a string seventeen characters long, even though the longest common subsequence is higher with the seventeen-character string (three versus one). The corrected Gestalt pattern matching algorithm produces similarity scores that are less influenced by length discrepancies and more influenced by the longest common subsequence.

While the Gestalt pattern matching algorithm works well for the comparison of words in a text document that tend to be roughly the same length, the algorithm works less well for the comparison of molecular pathways that can differ dramatically in length. For the latter, the algorithm tends to generate similarity scores that are dominated by pathway length and not the longest common subsequence. To avoid this potential artifact, we added a minimally perturbing correction term to the denominator of Equation 1:

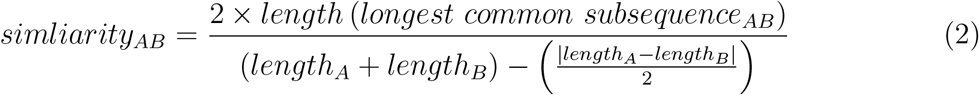

This correction term subtracts half the length difference between the two pathways being compared from the combined length of both pathways, acting as a “penalty” towards the similarity score if the two pathways being compared are of drastically different lengths (Figure 2B). If the two pathways have similar lengths, the correction term becomes zero.

## LPATH application to an alanine dipeptide conformational transition

### Simulation details

Our benchmark application of LPATH involved the conformational transition of alanine dipeptide from the C7_eq_ to C7_ax_ states (Figure 3). Our simulations employed the AMBER ff14SBonlysc force field^23^ for alanine dipeptide and generalized Born implicit solvent.^24,25^ A 4-fs timestep was enabled in all simulations by using a hydrogen mass repartitioning scheme. WE simulations were run with a *τ* =100 ps and a two-dimensional progress coordinate consisting of *ϕ* and *ψ* backbone torsional angles. Fixed bins for WE were placed only along the *ψ* dimension of the progress coordinate at a 20° interval between 0° and 360°. Trajectory coordinates were saved every 4 ps. The five, independent WE simulations generated 80 successful pathways in 14.6 μs of aggregate simulation time. Each simulation was completed in 21 hours using 8 CPU cores of a 3.5 GHz Intel Xeon CPU in parallel. An equivalent 14.6 *μ*s of cMD simulation yielded only 30 successful pathways, which was not sufficient for a robust analysis of pathways. Nevertheless, we have included an example of how to analyze cMD simulations using LPATH in the example GitHub repository.

**Figure 3.**
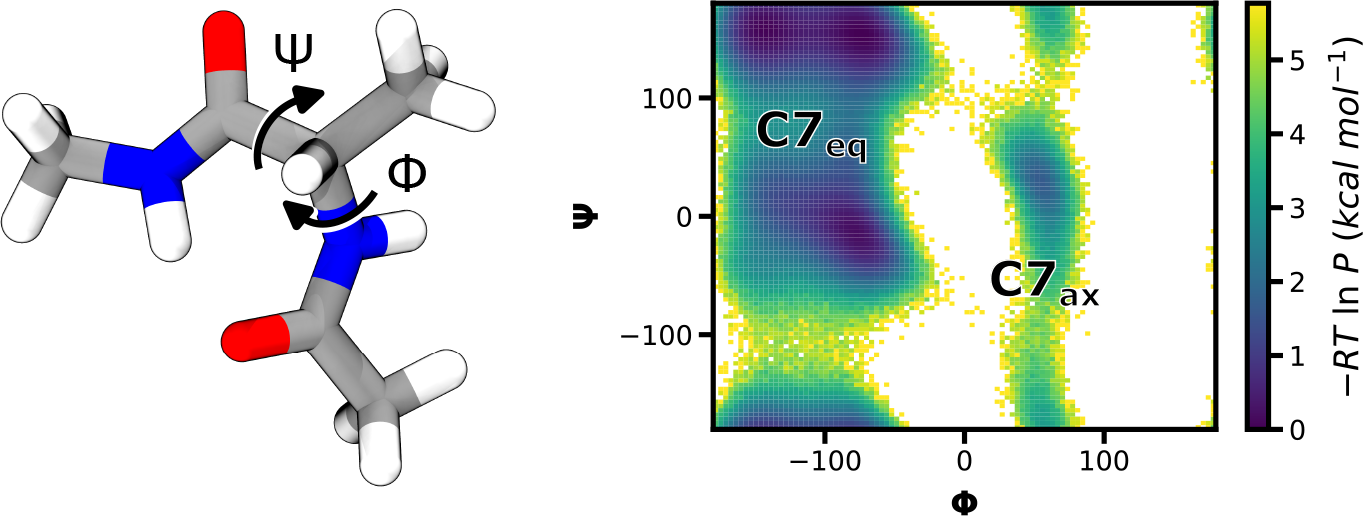
The C7_eq_ to C7_ax_ conformational transition of an alanine dipeptide as a benchmark application. Alanine dipeptide, capped with acetyl and N-methyl groups, is shown alongside a probability distribution of conformations as a function of the *ϕ* and *ψ* backbone torsional angles. In this work, we focus on the transition between C7_eq_ and C7_ax_, which involves surmounting a relatively large energy barrier (∼5 kcal/mol) along *ϕ*.

We strongly recommend generating multiple independent simulations whenever possible to assess the variation between runs, as we have done in this work. To combine multiple WE simulations prior to application of the LPATH workflow, use the WESTPA’s `w_multi_west` tool with the `--ibstates` flag.

### Discretizing the configuration space via clustering

To discretize the configuration space, we first assigned the source and target states to the C7_eq_ and C7_ax_ states, respectively, of alanine dipeptide and then clustered conformations between the source and target states. To circumvent the memory costs of clustering a large number of conformations, we implemented and subsequently applied a two-stage approach that first trains a machine learning model on a subset of conformations that have been cluster-labeled with a clustering method and then uses the resulting model to predict cluster labels for the remainder of the dataset. The use of a pre-trained model in this step is optional and only used due to the large number of conformations that needed to be assigned states. In the first stage, we applied hierarchical agglomerative (bottom up) clustering with an “average” linkage criteria on a subset (data from the final 50 iterations of each WE simulation) of the conformations, tuning the distance threshold (i.e., to 75) to yield six clusters that correspond to known conformational states of alanine dipeptide in *ϕ*/*ψ* torsional angle space (Figure 4A). We then used the resulting cluster-labeled conformations to train a k-nearest neighbors classifier model (N=5) on the cluster-labeled subset of conformations. In the second stage, we used the classifier model to predict cluster labels for the remainder of the dataset. As validation of this two-stage approach, the centroids of the final clusters matched the centroids from the training set (Figure 4B). To avoid any artifacts due to the periodicity of the torsional space, we adjusted the range of *ϕ* angles from -180° to 180° to be between -210° and 150° prior to clustering the conformations.

**Figure 4.**
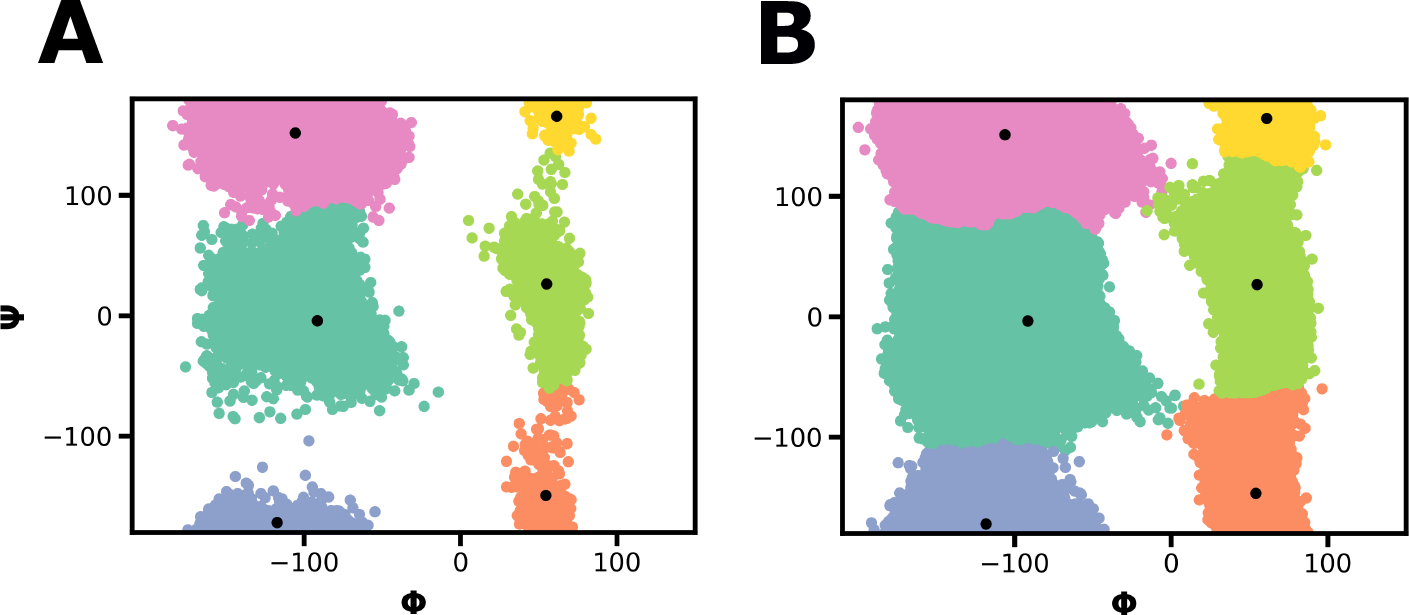
Validation of our two-stage clustering approach for a WE simulation of alanine dipeptide. A) Clusters resulting from hierarchical agglomerative clustering of a subset (every 4 ps from the final 50 iterations) of the WE simulation data. Each data point (conformation) is colored by cluster. Centroids of these initial clusters are indicated by black dots. B) Clusters assigned to the full simulation dataset based on a machine learning (k-nearest neighbors) classifier model that was trained on the cluster-labeled subset of data in A). The corresponding centroids are indicated as black dots and are in close agreement with those from the subset in A).

### Pathway clustering identifies two distinct pathway classes

After assigning states, we calculated the similarity score of each pair of pathways using our modified Gestalt pattern matching algorithm (Equation 2). Next, we clustered all successful pathways with hierarchical agglomerative clustering using the “Ward” linkage criteria and the distance (*d* = 1 −*similarity*_*AB*_) between each pair of sequences A and B. We then identified distinct pathway classes based on a dendrogram (tree diagram) of the clustering results. Figure 5A displays the dendrogram constructed using 80 successful pathways from the set of five, independent WE simulations. Each vertical “leaf” in the dendrogram represents a pathway, which connects to other pathways through horizontal “nodes”. Dendrogram branches with nodes that are similar to each other are closer together in the vertical direction. We identified the most distinct grouping of pathways into classes by positioning a horizontal line at a point that divides the dendrogram vertically between nodes with a maximum distance separation. For our WE simulation, a horizontal line at y=1.25 divides the dendrogram at the maximum vertical distance between nodes, identifying two pathway classes. We advise positioning this line at a few different positions and noting its impact on the number and features of the resulting pathway classes. Often, drawing the line too low in the dendrogram will generate pathway classes with redundant features (such as tracing the same pathway through phase space). If the number of pathway classes is unclear from the dendrogram, we recommend revisiting steps 1 and 2 of the LPATH workflow to ensure that (i) a minimum of three states was used for discretization of configuration space, (ii) at least 50 total pathways were extracted and (iii) each pathway contains at least 10 frames.

**Figure 5.**
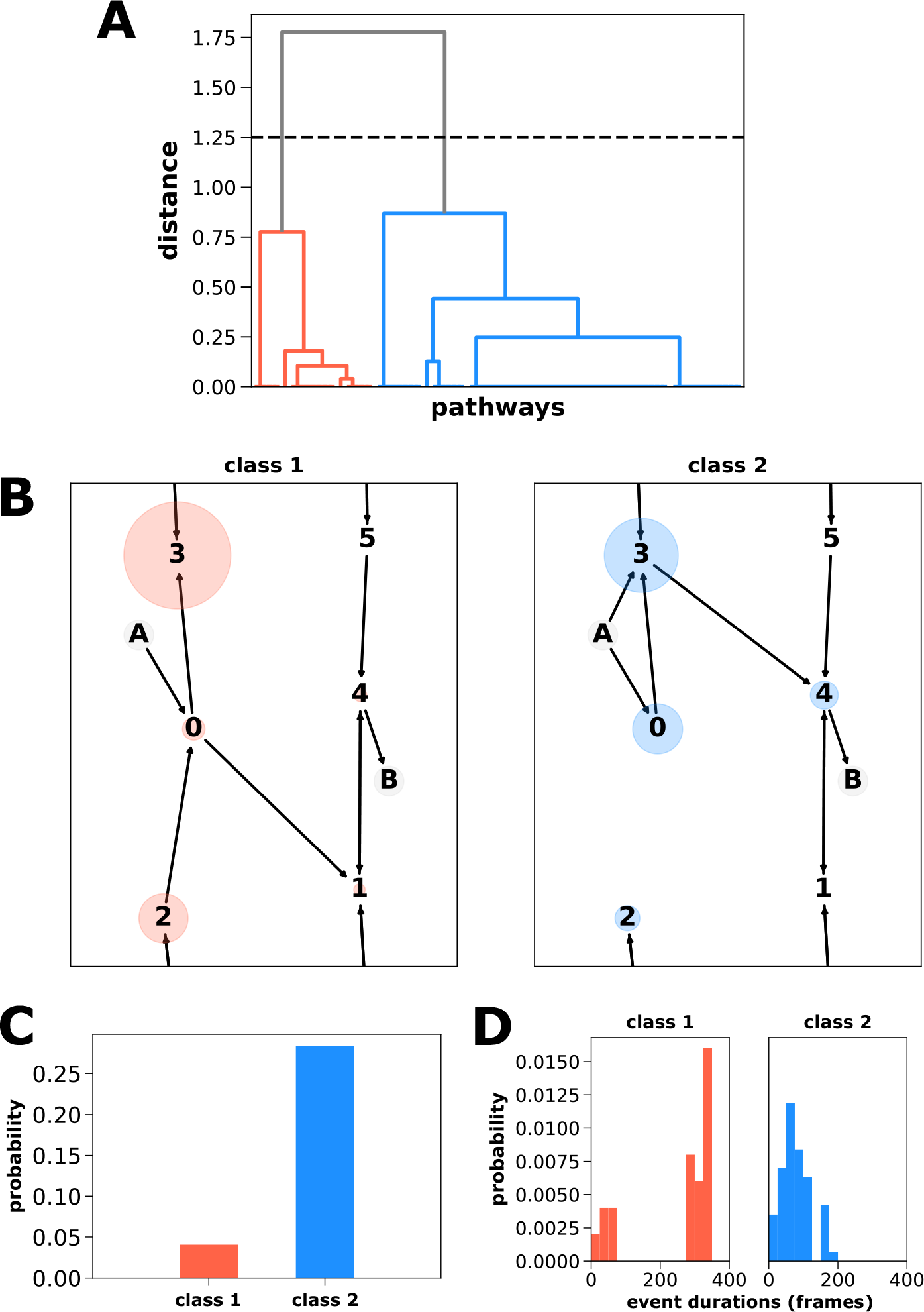
Analysis of pathways for the C7_eq_ to C7_ax_ transition of alanine dipeptide generated using five WE simulations with coordinates saved every 4 ps. A) Dendrograms of successful WE-generated pathways (N=80), reveal two distinct pathway classes 1 and 2 based on the cluster distance indicated by the horizontal dashed line. B) Directed network plots reveal that pathway class 1 involves an upper route from state 0 to state 1 and pathway class 2 involves a lower route from state 3 to state 4. C) Histogram of the two pathway classes indicating that the upper route (blue) is more probable than the lower route (red). D) Histograms of the event duration (barrier crossing) times for each pathway class indicate that the pathway classes are not determined solely based on pathway length, and that the upper, more probable route is more direct. This simulation dataset consisted of 5 WE simulations totaling 14.64 *μ*s of aggregate simulation time.

To determine how the two pathway classes differ in the mechanism of the C7_eq_ to C7_ax_ transition, we generated directional network plots of the fundamental “condensed” pathway routes through *ϕ*/*ψ* torsional space. (Figure 5B). Each node in the network plot corresponds to a defined state visited by the trajectories and is scaled according to the trajectory weights from the WE simulations. The C7_eq_ to C7_ax_ transition of alanine dipeptide appears to cross the main energy barrier in the *ϕ* dimension along two main “routes,” one from state 0 to state 1 (the lower route) and one from state 3 to state 4 (the upper route). A histogram of the two pathway classes (Figure 5C) reveals that the upper route is more probable (87.5%) than the lower route (12.5%). Based on the distribution of event duration (barrier crossing) times for each pathway class (Figure 5D), it is clear that both short and long pathways are grouped into the same classes, indicating that our modified Gestalt pattern matching worked as intended.

## Conclusions

The LPATH tool reveals distinct classes in the pathway ensemble by discretizing the configuration space into key states, extracting successful pathways and matching those pathways. The heart of the LPATH tool is the use of a custom score based on the Gestalt pattern matching algorithm from computational linguistics which clusters solely based on the matching of text strings representing the pathways. The generality of the pattern matching algorithm, which supports matching pathways of variable lengths, allows for a semi-automated workflow. We demonstrate the effectiveness of the LPATH tool in analyzing the pathway ensembles of alanine dipeptide from five independent WE simulations. The two distinct pathway classes identified by our tool correspond to “upper” and “lower” routes from the C7_eq_ to C7_ax_ conformational states. The interoperability of the LPATH tool enables straightforward implementation of alternate methods such as the geometric matching used in the PSA method and the Voronoi clustering used in the PHAT method.

## Notes

LTC serves on the scientific advisory board of OpenEye Scientific Software and is an Open Science Fellow with Psivant Sciences.

### Data and Software Availability Statement

Pathway analysis was performed with the open-source LPATH software package that is available on GitHub: https://github.com/chonglab-pitt/LPATH and deposited under DOI: 10.5281/zenodo.8403685. The open-source WESTPA software package will also be needed for some of LPATH’s functionality and is available on GitHub: https://github.com/westpa/westpa. The WESTPA HDF5 data file needed to reproduce these results, along with the associated plotting scripts can be found in the LPATH GitHub repository under the examples folder. Full documentation for using LPATH, including information needed to reproduce the results of this study, can be found here: https://lpath.readthedocs.io.

## Acknowledgement

We thank Daniel Zuckerman at Oregon Health & Science University for helpful discussions. We also thank Marion Silvestrini for coming up with the Pittsburgh/Plattsburgh example in Figure 2.

